# When Yeast Cells Change their Mind: Cell Cycle “Start” is Reversible under Starvation

**DOI:** 10.1101/2021.10.31.466668

**Authors:** Deniz Irvali, Fabian P. Schlottmann, Prathibha Muralidhara, Iliya Nadelson, N. Ezgi Wood, Andreas Doncic, Jennifer C. Ewald

## Abstract

Eukaryotic cells decide in late G1 whether to commit to another round of genome duplication and division. This point of irreversible cell cycle commitment is a molecular switch termed “Restriction Point” in mammals and “Start” in budding yeast. At Start, yeast cells integrate multiple signals such as pheromones, osmolarity, and nutrients. If sufficient nutrients are lacking, cells will not pass Start. However, how the cells respond to nutrient depletion after they have made the Start decision, remains poorly understood.

Here, we analyze by live cell imaging how post-Start yeast cells respond to nutrient depletion. We monitor fluorescently labelled Whi5, the cell cycle inhibitor whose export from the nucleus determines Start. Surprisingly, we find that cells that have passed Start can *re*-import Whi5 back into the nucleus. This occurs when cells are faced with starvation up to 20 minutes after Start. In these cells, the positive feedback loop is interrupted, Whi5 re-binds DNA, and CDK activation occurs a second time once nutrients are replenished. Cells which re-import Whi5 also become sensitive to mating pheromone again, and thus behave like pre-Start cells. In summary, we show that upon starvation the commitment decision at Start can be reversed. We therefore propose that in yeast, as has been suggested for mammalian cells, cell cycle commitment is a multi-step process, where irreversibility in face of nutrient signaling is only reached approximately 20 minutes after CDK activation at Start.

## Introduction

Cells must coordinate growth and cell division with metabolism to proliferate and to maintain cellular homeostasis. This holds particularly true for single cell organisms like budding yeast, that are exposed to constantly changing nutrient supply [1]. In yeast, nutrient supply directly affects metabolic rates and cell growth. Also cell size, and entry and progression through the cell division cycle are regulated in response to nutrients [2–5]. However, how the cell division cycle is responding to nutrients and which metabolic regulators interact with which components of the cell cycle machinery is still poorly understood [4].

One of the most critical points of cell cycle signaling is the commitment point at the end of G1, termed “Restriction point” in mammals and “Start” in yeast [6–8]. While the proteins involved in this decision have diverged, the architecture of cell cycle commitment is conserved from human to budding yeast [6, 9]. In yeast, the cell cycle inhibitor Whi5 inhibits the transcription factor SBF [10] (Figure 1A). This inhibition is partially relieved though dilution by growth and activation of SBF-dependent transcription by Cln3 [11–13]. The partial activation of SBF leads to the transcription of the activators of CDK, the cyclins Cln1 and Cln2. The Cln1/2-CDK complex hyperphosphorylates Whi5, which leads to its export from the nucleus and alleviation of SBF inhibition [12, 14]. This positive feedback loop leads to full SBF activation, and the G1/S transition can proceed [15]. The point where 50% of Whi5 has exited the nucleus is when the positive feedback loop becomes self-sustaining. This is considered the point of irreversible cell cycle commitment, Start [16–18].

**Figure 1:**
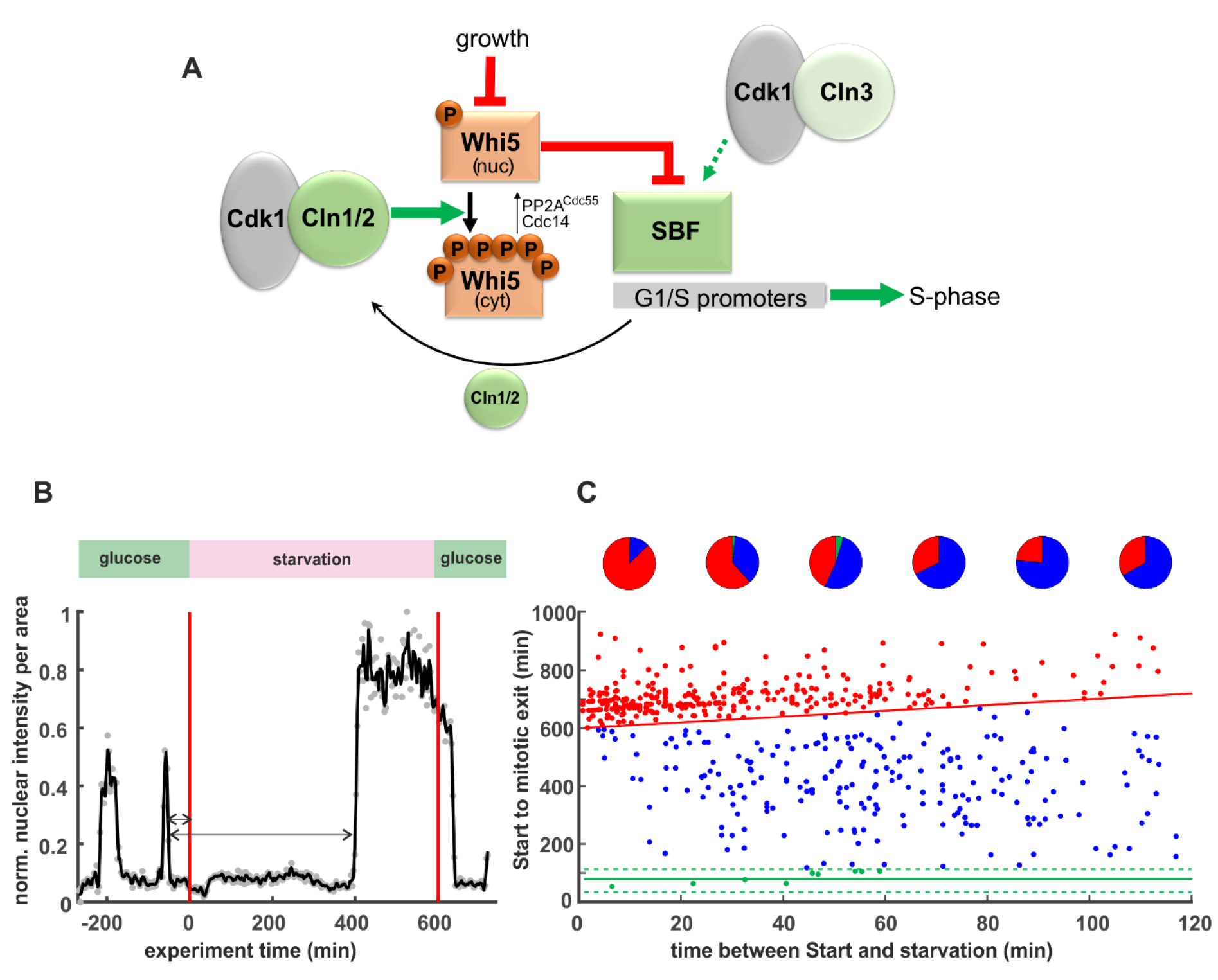
Post-Start cells respond to starvation. **A.** Model of Start based on [11, 12, 15, 17]. Cln1/2-CDK hyperphosphorylated Whi5 which leads to nuclear export **B.** Example trace of Whi5-mCherry single yeast cell in a starvation experiment. Cells were grown for eight hours on glucose minimal medium, ten hours in starvation medium (1% sorbitol minimal medium), then four hours on glucose minimal medium. Red lines indicate times of media switch. Grey dots represent original data, the black line represents data smoothed with a Savitzky-Golay filter. The arrows indicate the times used for the x and y axis in graph C. **C.** Cell cycle durations of single cells exposed to starvation. Cells were grown as described in B. Each dot depicts a single cell (n=579, from four independent experiments). The red line depicts the end of the 10-hour starvation period. Cells above this line continue their cycle only after glucose replenishment and are thus categorized as “permanently arrested”. The green lines depict the average duration in unperturbed cells +/- two standard deviations. Cells between the green and the red line are categorized as “delayed” cell cycle (such as the example cell in B). Pie charts depict the fractions of “permanently arrested” (red), “delayed” (blue), and “normal duration” (green) in 20-minute bins.

Leading up to Start, the cell integrates multiple internal and external signals - such as hormones and growth factors, stress, and cell size - to make the decision to commit to another round of DNA duplication and division [5–7]. It is well established that also nutrient signaling can promote or prevent Start [3, 4, 18, 19]. Cells that do not have a sufficient supply of all essential nutrients, will not pass Start and remain arrested in G1. However, what happens if essential nutrients are depleted *after* cells have passed Start, is largely unknown [4].

Evidence from our work and others suggests that cells can respond to nutrients in all phases of the cell cycle. For example, S-phase cells exposed to stress or a drop in glucose supply will transiently arrest the replication machinery by an inhibitory phosphorylation on Mrc1 [20]. Wood et al. found in single cell experiments that cells can enter a quiescence-like state outside of G1 when responding to acute starvation [21], in agreement with earlier population-based studies showing that even budded cells can enter quiescence [22]. A recent report also demonstrated that cells can arrest their cell cycle in a “high CDK-state” [23] when facing nutrient perturbations. Despite these individual examples, a comprehensive picture of how cells proceeding through the cell cycle respond to nutrient signals is still lacking. Specifically, we do not know which cell cycle regulators receive nutrient-dependent signals, or what phases of the cell cycle cells can delay or can stably arrest to wait for an improvement of nutrient supply.

Here, we investigated the response of post-Start cells to acute carbon starvation. We analyzed thousands of single cells by monitoring Whi5 and other cell cycle regulators during live cell imaging. We found that cells stably arrest their cell cycle in G1 or G2 phases, while S-phase seems to be only transiently delayed. Surprisingly, cells that had already passed Start, could translocate Whi5 back into the nucleus when starved within the first 20 minutes post-Start. This nuclear re-entry of Whi5 corresponded to a reversal of Cdk1 activation and cells became sensitive to mating pheromone again.

We conclude that Start remains reversible in response to nutrient perturbations for approximately 15-20 minutes. Thus, the current model of Start [6, 7, 16, 17, 24] is incomplete and Cdk1-Cln1/2 activation is not the final point of cell cycle commitment.

## Results

The impact of nutrient signaling on cell cycle progression downstream of Start is still largely unknown. Here, we investigated how acute carbon starvation impacts the yeast cell cycle using live cell imaging and microfluidics cultivation. Importantly, we worked with prototrophic cells on glucose minimal medium without amino acids, to ensure cells do not have an alternative carbon source. When starving the cells, we switched to medium containing 1% sorbitol, which is not metabolized by cells, but osmotically balances the medium. With this setup, we are confident to insulate specifically the effect of carbon supply and signaling. In our typical carbon starvation experiments, we first grew cells for several hours on glucose medium to adapt cells to the microfluidic environment and obtain data on the unperturbed cell cycle. Cells were then switched to starvation medium for ten hours (unless stated otherwise), after which glucose was replenished for at least four hours.

Cells in the microfluidics cultivation platform grow asynchronously, therefore we could observe cells in all phases of the cell cycle when the medium was switched to starvation conditions. Out of all cells, we specifically analyzed single cells that had already passed Start when they were exposed to carbon depletion. Start was determined by monitoring endogenously fluorescently tagged Whi5, whose exit from the nucleus marks the Start transition [17, 18, 25]. We determined the time that had passed between Start and when a cell was faced with carbon depletion (Figure 1B, x-axis on Figure 1C), and the time that these cells spent from Start to the end of mitosis, as monitored by the re-entry of Whi5 into mother and daughters (Figure 1C, y-axis). We analyzed over 500 post-Start cells (from four independent biological experiments) and found three different types of responses: A small fraction (<2%) of cells completed their cell cycle in a similar time as non-starved cells (Figure 1C, green dots), 39 % of cells completed their cell cycle, but with a much longer duration than non-starved cells (Figure 1C, blue dots, we refer to these as “delayed cell cycle”), and 59 % permanently arrested their cell cycle for the entire ten hours of the starvation period (Figure 1C, red dots). This arrest was typically stable even when the starvation period was extended to 16 hours. Independent of their cell cycle or arrest state, almost all cells continued normal cell cycles after glucose was replenished, indicating that cells do not lose viability or proliferative capacity during long-term starvation-induced arrest. The fraction of cells that consecutively stopped proliferation was below 1% in our assays, but this fraction gradually increased if the duration of starvation was extended.

Our data confirm recent studies that cells respond to nutrient depletion in all phases of the cell cycle [21, 23], however the mechanisms behind the cell cycle delay and arrest have not been determined. We decided to focus on those cells that enter a stable arrest to explore the mechanisms leading to and stabilizing the arrest. When analyzing arrested cells, we noticed a surprising phenomenon: Cells that were exposed to starvation could translocate the Start inhibitor Whi5 back into the nucleus (Figure 2, Movie 1, Figure 3A). Given this surprising result, we wanted to confirm that Whi5 exit in these cells indeed corresponded to the CDK positive feedback loop being activated and thus cells actually passing Start. We monitored Cln2 promoter activity using a fluorescent protein driven by the Cln2 promoter, but realized that the inherent delay caused by fluorescent maturation hindered sufficient time resolution. To increase time resolution for analyzing the promoter activity, we turned to the dPSTR system (developed by the Pelet lab [26]) which turns a transcriptional signal into a localization change of a fluorescent reporter (see also Supplementary Figure 1). With the Cln2-dPSTR reporter we indeed showed that all analyzed cells activated Cln2 promoters at Whi5 exit (example cell in Figure 2C), and thus had passed *Start* (as defined in the textbooks) when translocating Whi5 back into the nucleus a few minutes after being faced with starvation. This nuclear translocation of Whi5 after Start cannot be explained by the current model of Start, which states that Whi5 exit leads to *irreversible* commitment to cell cycle progression. However, Whi5 re-entries have been noted, but not explained, in several previous studies [21, 25, 27, 28]. We thus decided to examine when, how and why Whi5 re-entry occurs.

**Figure 2:**
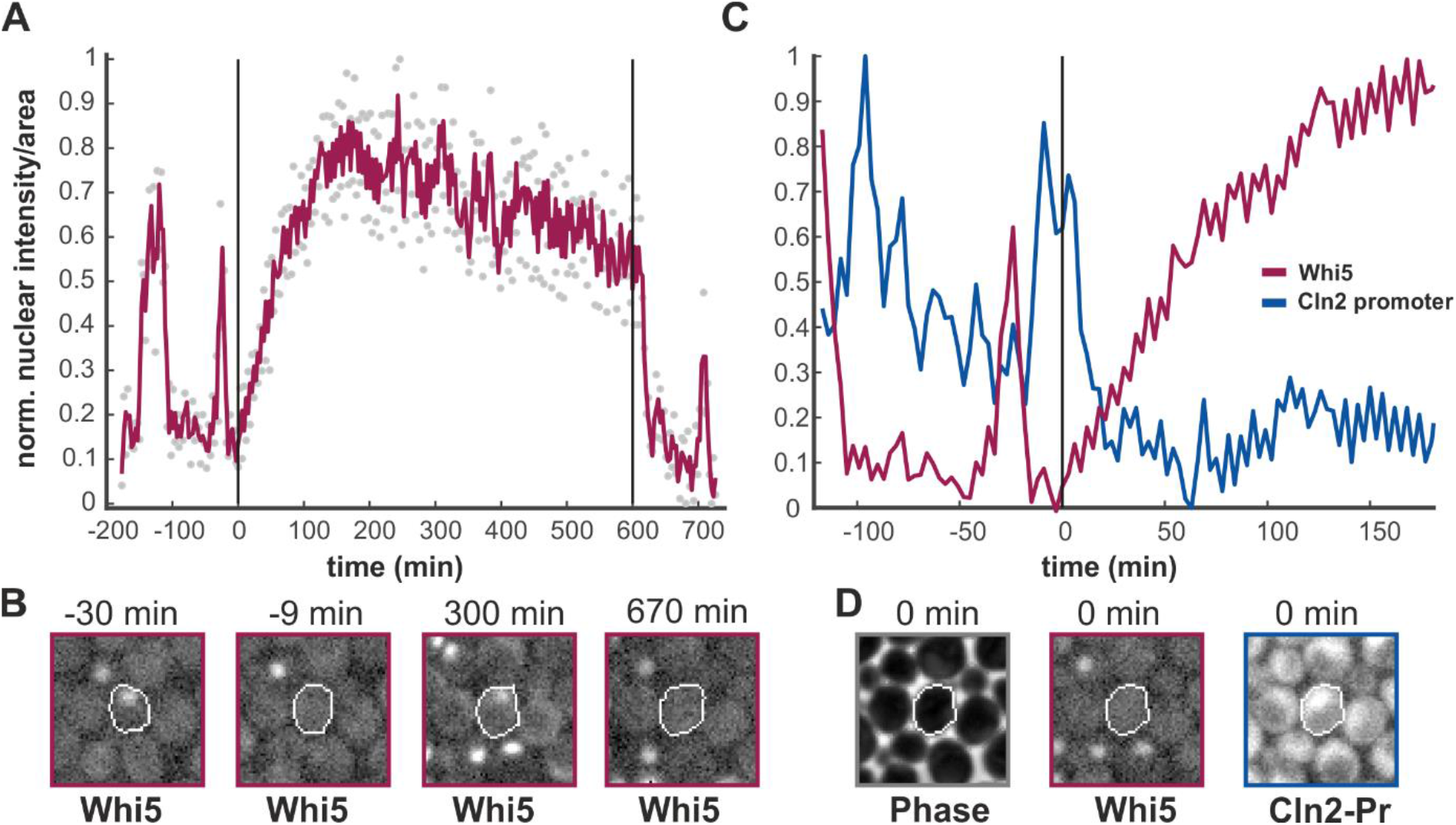
Whi5 re-enters the nucleus upon starvation. **A.** Whi5-mCherry trace of an example cell that re-imports Whi5 upon starvation. Grey dots represent raw data, the magenta line results from a Savitzky-Golay smooth. Starvation was induced at t=0 minutes and relieved after 10 hours as indicated by the vertical black lines. **B.** Images of the cell trace shown in A at indicated time points. **C.** Excerpt of the Whi5-trace shown in A including data from a Cln2-dPSTR expression reporter (blue line). Nuclear localization of the reporter indicates activity of the Cln2 promoter. **D.** Images from the cell shown in C at the time point immediately after starvation is induced. Absence of Whi5 and presence of the Cln2 reporter in the nucleus indicate that this cell had passed Start when it was exposed to starvation. See also Movie 1.

**Figure 3:**
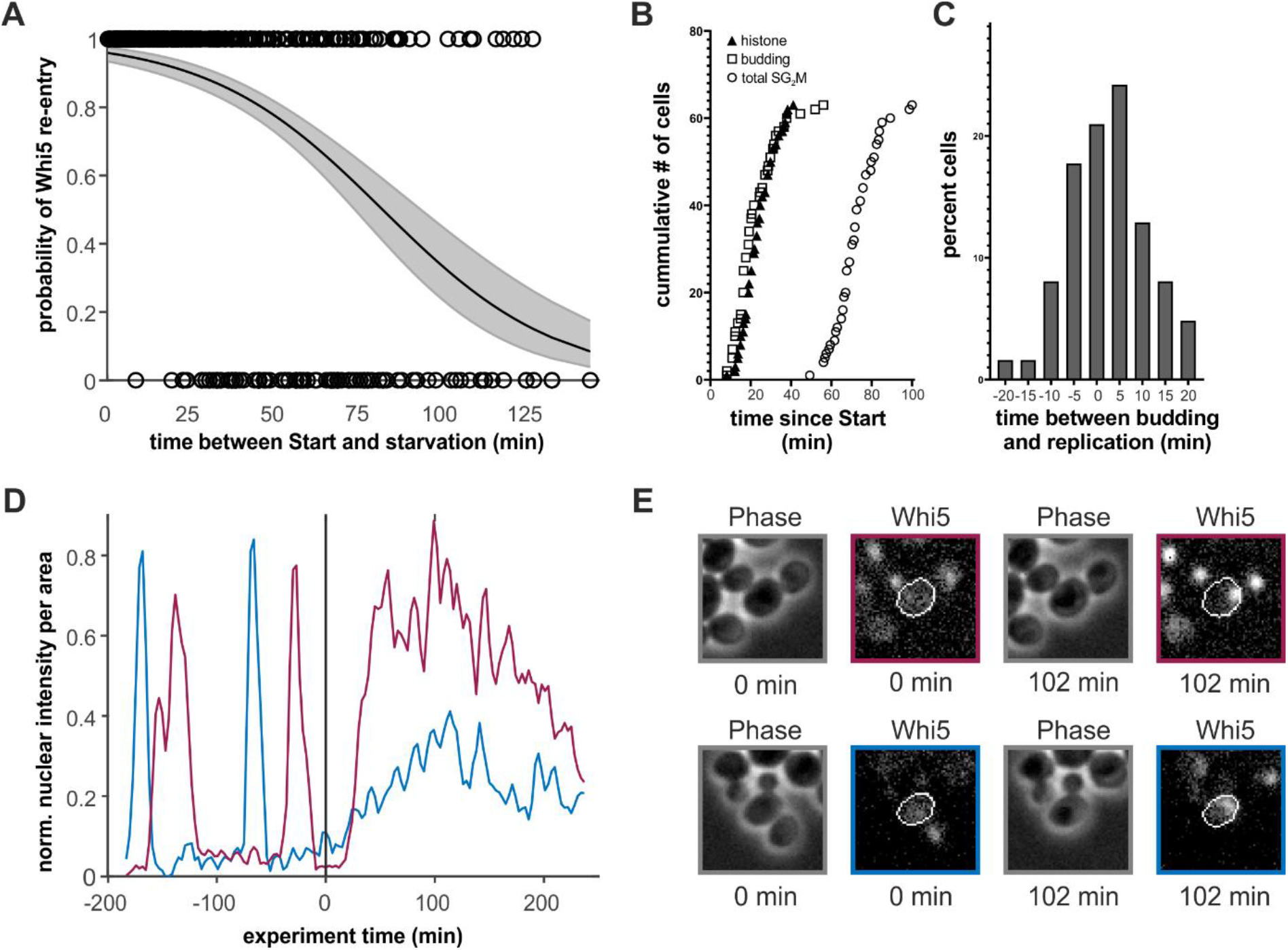
Early and late Whi5 re-entries. **A.** The probability of Whi5 nuclear re-entry after starvation. Black circles indicate single cells (n=849, from 5 independent experiments) that translocate Whi5 back into the nucleus. The x-axis denotes the time that had passed since Whi5 exit until the medium was switched to starvation medium. The black line indicates a logistic regression of the single cell data, where the grey area denotes the 95% confidence intervals. **B.** Timing of budding and replication in relation to Start of cells growing on glucose minimal medium. Htb2-TFP fluorescence increase was used as a proxy for replication onset. 65 single cells growing on glucose minimal medium were observed and the time between Start, budding, and onset of histone production were recorded. S/G2/M corresponds to the time between Whi5 exit, and Whi5 entry in mother and bud at the end of mitosis. **C.** Histogram of the time difference between budding and increase of Htb2 fluorescence in the 65 cells described in B. **D.** Example traces of Whi5 of two cells exposed to starvation at t=0 min. The magenta line indicates a cell with a fast and steep re-entry, the blue line indicates a cell with a slow re-entry. **E.** Images of the two cells described in D at the indicated timepoints.

Based on >800 observed cells, we calculated the probability of Whi5 re-entry, given the time between Start and the exposure to starvation (Figure 3A). While the probability was highest in the first ten minutes, the probability for re-entries gradually dropped, but never reached zero. This seemed surprising given that cells begin replicating their DNA at approximately 22 minutes after Start (Figure 3B), so we did not understand the functional relevance of these late re-entries. In this context however, we noticed that there seemed to be two qualitatively distinct types of Whi5 re-entries. Cells that were early in the cell cycle showed an immediate and steep Whi5 re-entry. In contrast, cells that were later in the cell cycle and had a large bud showed a slow Whi5 re-entry (example cells in Figure 3D and E). We hypothesized that these early and late nuclear re-entries of Whi5 were mechanistically and functionally distinct.

We thus looked more closely at the dynamics of Whi5 re-entry. We calculated the re-entry slopes and determined the budding state of these cells (Figure 4A). Confirming our initial observation, the median slope of unbudded cells was approximately 2.5-fold higher than in budded cells. However, there were some cells with small buds with very steep Whi5 re-entries, and on the other hand cells with no obvious buds that showed slow re-entries like budded cells. Thus, the budding event per se is not what distinguishes the two different types of Whi5 behavior.

**Figure 4:**
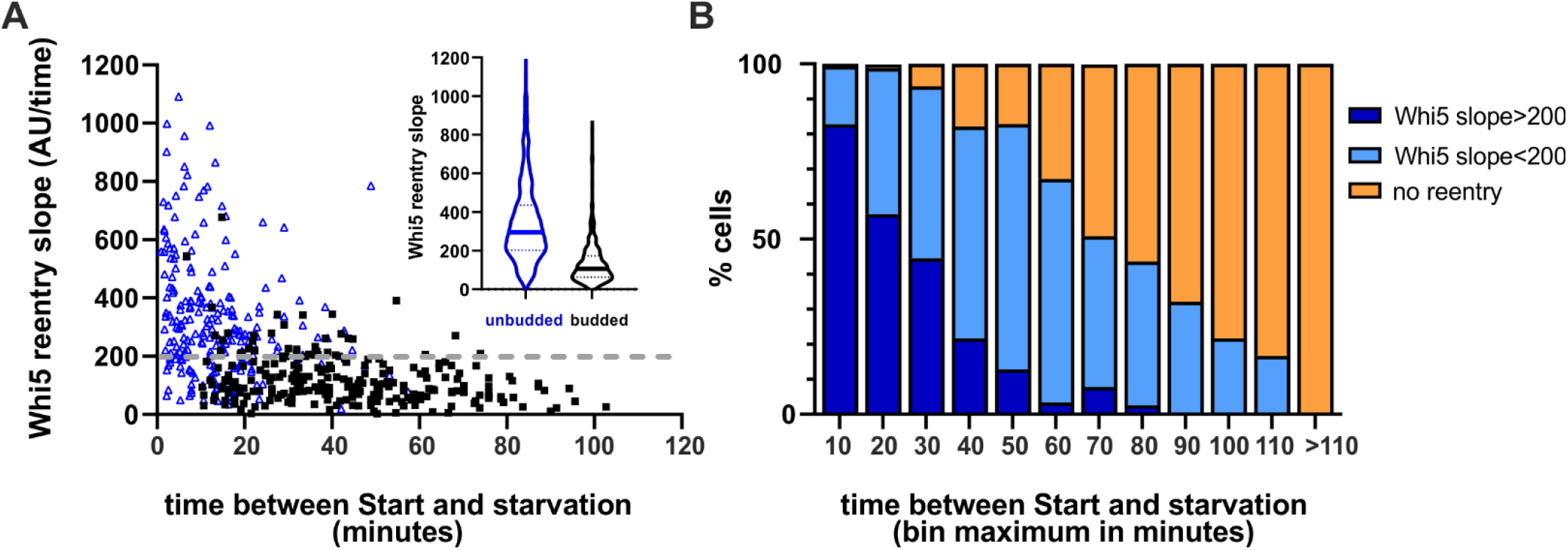
Two different types of Whi5 re-entries. **A.** We determined the slope of Whi5 nuclear re-entry (regression of the first 30 minutes of nuclear Whi5 increase) in budded (black squares) and unbudded cells (blue triangle). We analyzed 680 cells from 5 replicate experiments. We picked 200 AU/time (dotted grey line) as the threshold between fast and slow re-entries. Inset shows a violine plot of the slopes of all budded (black) and unbudded (blue) cells, where the solid lines depict the median of the distribution. **B.** Distribution of fast, slow, and no re-entries depending on how long after start cells experience starvation. (849 cells from 5 replicate experiments).

Budding typically co-occurs with the onset of replication, but is not mechanistically coupled (Figure 3C) [29]. We thus hypothesized that the difference between Whi5 behavior is due to the onset of replication, i. e. the actual beginning of S-phase. There is so far no canonical marker for observing S-phase initiation in budding yeast; typically, the increase in histone concentration is used. Unfortunately, upon starvation, the fluorescence from TFP- or mCherry-labelled histones increased in all phases of the cell cycle likely due to sensitivity of the fluorophores to changes in the cellular environment. We therefore could not precisely determine the onset of histone production in our starved cells. However, the timing of histone production onset in unperturbed cells (Figure 3B) corresponds well to the timing of the change in Whi5 behavior (Figure 4A). We therefore decided to group the cells according to their slope of Whi5 re-entry, rather than by budding. We chose a cut-off of 200 arbitrary intensity units/min (dotted grey line in Figure 4A) to separate “fast” re-entries, which we interpret as pre-replication cells, and “slow re-entries” which we interpret as cells that have started replication and eventually arrest in G2.

Neither the fast nor the slow re-entries can be explained with our current understanding of Whi5 and Start regulation. However, since a function of Whi5 outside of Start is not known, we decided to focus on those cells re-importing Whi5 at the G1/S transition. Fast Whi5 re-entries were very commonly observed: Within the first 20 minutes after Start, over 70% of cells translocated Whi5 back into the nucleus with a steep slope (Figure 4B); in the first 5 minutes after Start 99% of cells re-imported Whi5. We hypothesized that this Whi5 re-entry corresponds to a decrease in Cln1/2-CDK activity. If Whi5 re-entry indeed corresponds to a decrease in Cln1/2-CDK activity, the cells would have to go through another round of the positive feedback loop of Whi5 exit and Cln1/2 activation after glucose replenishment. We therefore used the Cln2 promoter constructs to see what happens in these cells after glucose replenishment. Indeed, cells that had translocated Whi5 back at the onset of starvation, later showed another spike in Cln2 promoter activity once glucose was replenished, concurrent with Whi5 exit (Figure 5 A-C). Notably, those cells classified as “slow re-entries” (Whi5 re-entry slope <200 AU/min) did not show a second round of Cln2 promoter activity before progressing with their cell cycle (Figure 5C). This confirms that these cells had arrested in S or G2, and no longer required Cln1/2 activity to continue their cell cycle.

**Figure 5:**
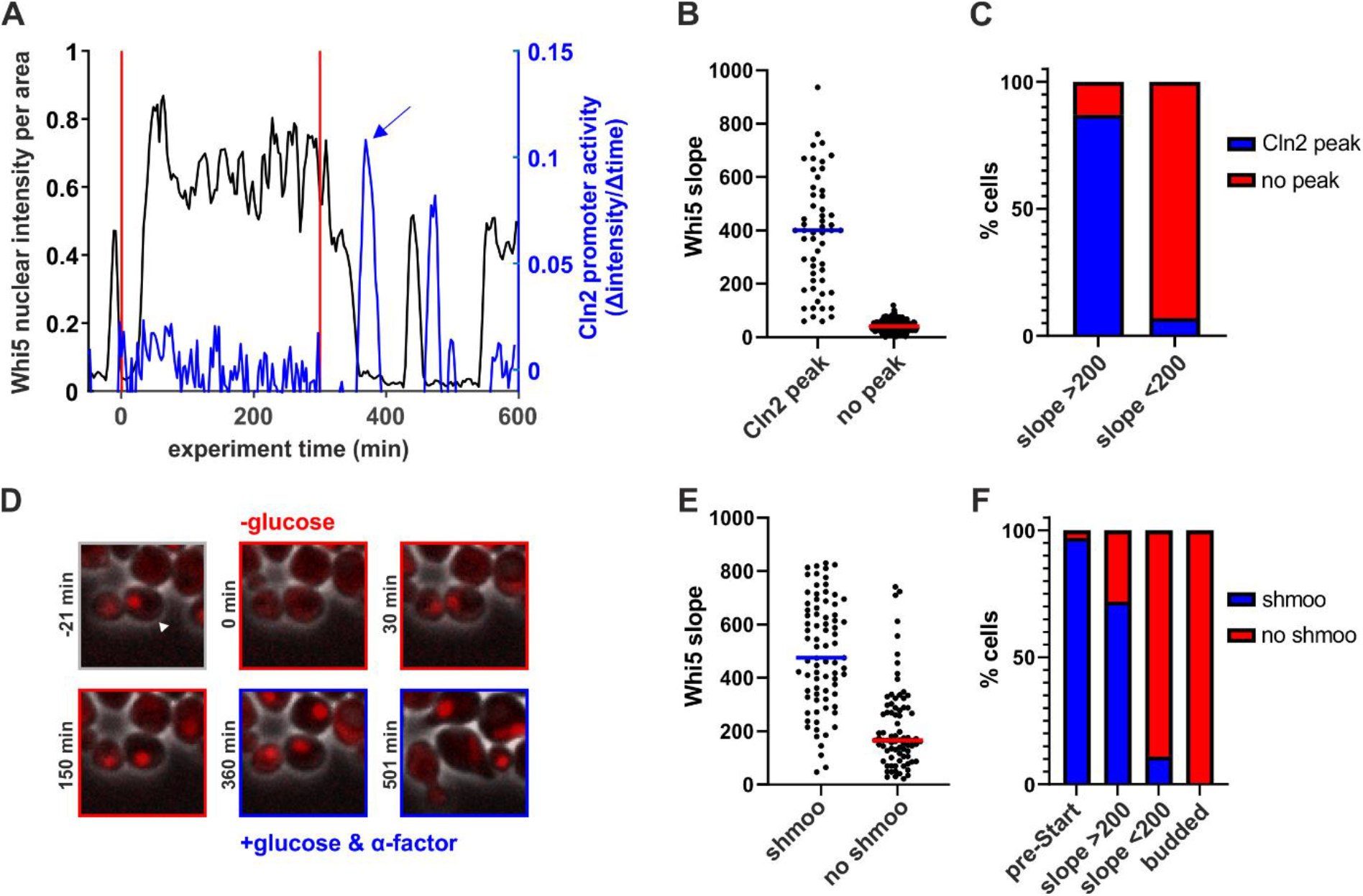
Early Whi5 re-entries lead to functional Start reversal. **A.** Example cell showing that the Cln2 promoter (blue) fires upon glucose replenishment after Whi5 re-entry (black). Vertical red lines indicate beginning and end of the starvation phase. The promoter activity was approximated by the change in fluorescence of a Cln2-Promoter-Neongreen construct (see also Supplementary Figure 1). **B** Whi5 re-entry slopes of cells with and without Cln2 promoter activity after glucose replenishment. (153 cells from 3 replicate experiments). **C**. Percentage of cells that show Cln2 promoter activity after glucose replenishment in cells that re-imported Whi5 with slopes above or below our threshold of 200 (Figure 4). Cells without Whi5 re-entry never showed a Cln2 peak after glucose replenishment. **D.** Example cell that re-imports Whi5 after starvation and responds to alpha-factor after glucose replenishment. **E.** Whi5 re-entry slopes of cells that shmoo or not shmoo after glucose replenishment and alpha-factor addition. **F.** Percentage of cells that respond to alpha-factor addition by shmooing threshold of 200 (Figure 4). The first bar describes cells that were arrested in a normal pre-Start G1. The two middle bars include cells that exported and re-imported Whi5 with no visible buds. The right bar includes all cells that were budded at the time of starvation and glucose replenishment. See also Movie 2.

These data on Whi5 localization and Cln2 promoter activity strongly suggested that activation of CDK at Start -supposedly the point of *irreversible commitment*- is in fact *reversible* under starvation. To test whether cells with Whi5 re-entries were functionally truly reversing cell cycle commitment, we tested their sensitivity to mating pheromone. We starved cells and then exposed them to the mating pheromone alpha-factor at the same time when glucose was replenished. Most cells with Whi5 re-entries during starvation later responded to alpha-factor by shmooing, just like cells that were in a normal pre-Start G1 state (Figure 5D-F, Movie 2), albeit the fraction of shmooing cells was lower. This sensitivity to mating pheromone confirms that the positive feedback loop defining “Start” can be reversed within the first ~20 minutes. Thus, the current model of irreversible cell cycle commitment does not hold true vis-à-vis nutrient perturbations.

Having established the functional reversibility of the Whi5-CDK positive feed-back loop, we next wanted to understand the mechanism by which the G1/S transition is interrupted and Whi5 is translocated back to the nucleus. One plausible explanation is that the expression of the G1 cyclins is targeted by metabolic regulators to interrupt the G1/S transition. We thus investigated the possible role of known transcription factors involved in starvation responses and quiescence: The starvation responsive transcription factors Msn2, Msn4, Xbp1 [30], have all been shown to bind the Cln1/2 promoters (data summarized on yeastract [31]); and the transcription factors Msa1 and Msa2 are important for arresting cells in G1 in preparation of quiescence [32]. While these transcription factors are clearly relevant for long-term starvation responses, they do not seem to be essential in driving Whi5 re-entry during acute starvation (Figure 6). The *xbp1, msa1msa2*, and *msn2msn4* deletion mutants all still translocated Whi5 back into the nucleus. We note however, that the re-entry dynamics of Whi5 appear different in the *msn2msn4* mutant compared to wildtype. The distinction between early and late Whi5 re-entries based on the slope of Whi5 re-import did not hold for all mutant cells. We observed both early re-entries which were slow and late re-entries with a very steep slope, for which we currently cannot offer a straight-forward interpretation. Thus, while the initial decision to interrupt Start is probably not caused by these transcription factors, they likely contribute (directly or indirectly) to full Start reversal and the stabilization of the G1 arrest.

**Figure 6:**
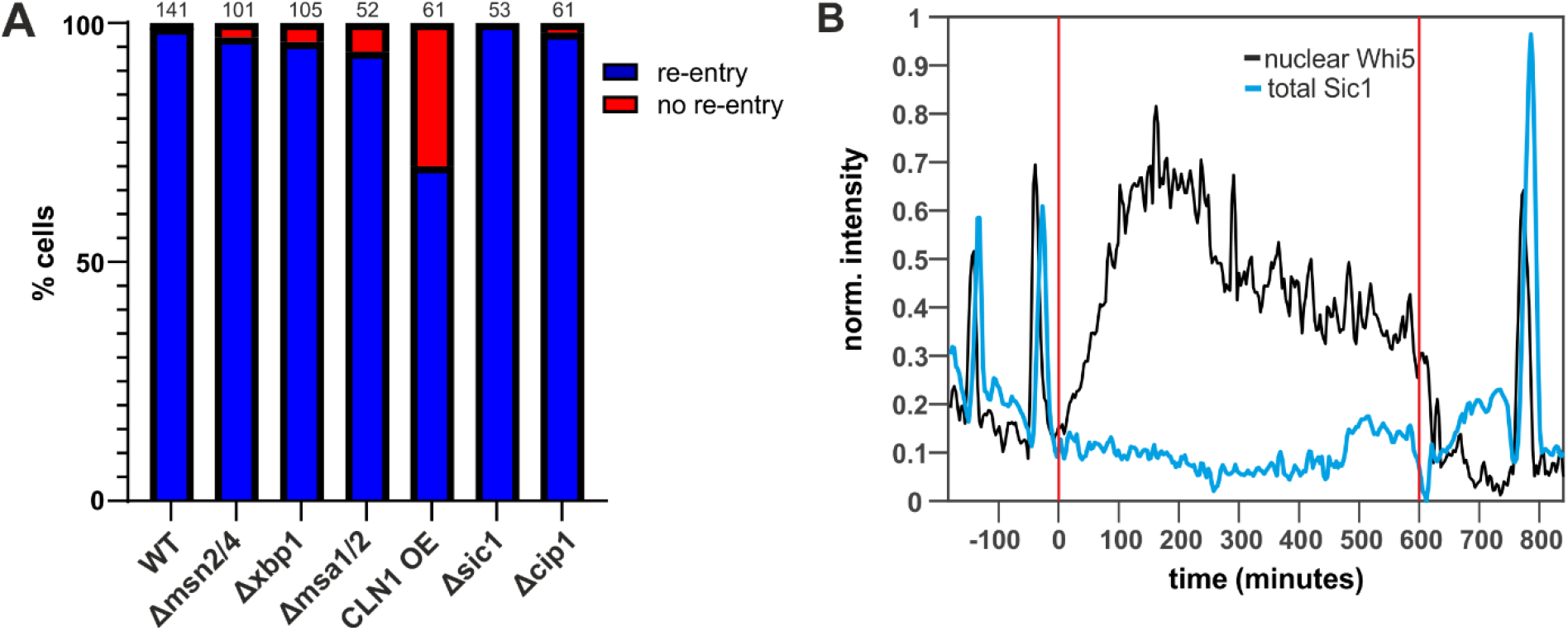
Deletion mutants still show Whi5 re-entries. **A.** We constructed the indicated mutants and monitored their Whi5 response to starvation. Depicted is the fraction of cells that re-imports Whi5 when starved within 15 minutes after Start. Numbers over the bars indicate total number of cells that were analyzed, cells were from at least three independent biological replicates. **B** Example of a cell where Whi5 (black) re-enters the nucleus upon starvation despite completed Sic1 (blue) degradation.

To test whether another transcription factor repressing cyclin expression may be essential for initiating Whi5 re-entry, we expressed Cln1 constitutively from a strong synthetic promoter, which is even active under starvation (Δcln1Δcln2Δcln3, estradiol-inducible-Cln1, from [33–35], see also Supplementary Figure 2). Even in these cells overexpressing the activating cyclin, Whi5 still re-entered the nucleus upon starvation in over 70% of the cells in the first 15 minutes (Figure 6). The 30 % reduction in cells re-entering Whi5 is presumably caused by an overall acceleration of the G1/S transition due to high Cln1 levels. Thus, while repression of G1 cyclin expression is likely important for establishing and stabilizing quiescence, it is not essential for reversing Start under acute starvation.

Since Cln1/2 repression does not seem to be the initial target for interrupting Start, we wondered if the downstream inhibitor Sic1 could play a role. Sic1 is a well-established inhibitor of CDK-cyclin complexes and needs to be degraded to transition to S-phase. Sic1 has been shown to be stabilized by Hog1 phosphorylation during the hyperosmolarity response [36, 37] and therefore seemed like an obvious target also for nutrient signaling. We thus fluorescently labeled Sic1 and determined the amount of Sic1 present in cells that translocated Whi5 versus cells that did not. However, Sic1 concentrations (total or nuclear) were not predictive of whether a cell reverses Start (Figure 6B and Supplementary Figure 4). In agreement with this result, a deletion mutant of Sic1 did not lead to less Whi5 re-entries (Figure 6A). In fact, due to a slightly slower cell cycle, the Sic1 deletion mutant translocated Whi5 back to the nucleus for an even longer time after Start. Similarly, a deletion mutant of the p21-like inhibitor Cip1 [38] did not impact Whi5 translocations (Figure 6A).

We thus investigated whether Whi5 itself is the target of starvation signaling as it has been reported for high-osmolarity signaling [39]. Whi5 has 12 reported CDK phosphorylation sites, whose phosphorylation leads to Whi5 release from promoters and exit from the nucleus at Start [12]. Additionally, there are at least seven non-CDK sites, which have been confirmed by mutational analysis ([12] and Jan Skotheim, personal communication), but that have no clear function yet. We thus wondered if one of these 7 phosphorylation sites could be targeted to cause Whi5 nuclear re-import. We thus mutated all seven of these non-CDK phosphorylation sites on Whi5, but this did not prevent Whi5 re-entry upon starvation (Figure 7A).

**Figure 7:**
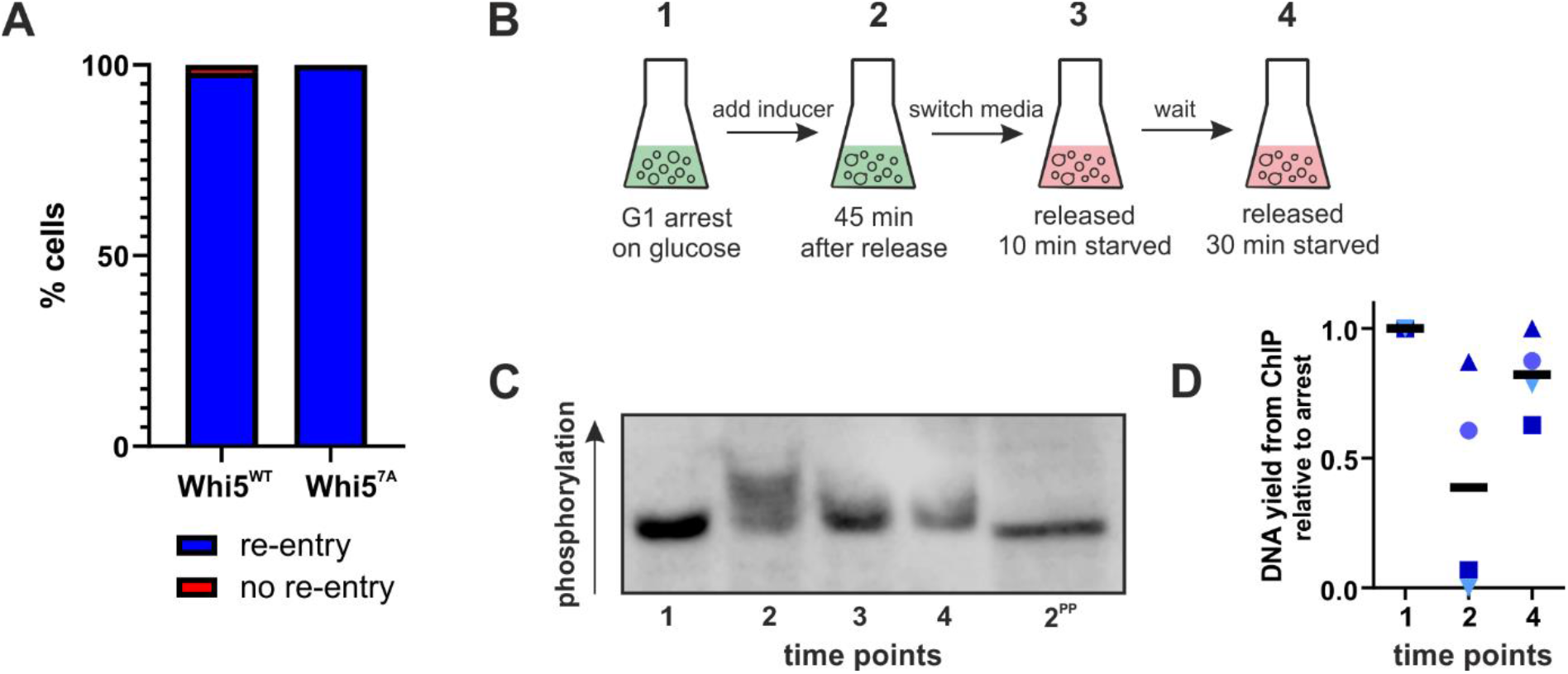
Whi5 phosphorylation during starvation. **A.** We introduced wildtype or the 7A phosphorylation site mutant (S78, S113, T114, S149, S276, T281, S288 mutated to alanine) Whi5 into a whi5-deletion strain and determined its response to starvation. Bars indicate the fraction of cells that re-import Whi5 when starved within 15 minutes after Start. **B.** Schematic of the experimental set-up used for B and C. **C.** Phostag-SDS-PAGE-Western blot of Whi5-V5. Numbers indicate the samples shown in B. 2^PP^ indicates sample 2 (cells growing on glucose, 45 minutes after a G1 release) treated with phosphatase. See Supplementary Figure 5 for a blot from a replicate experiment. **D.** Cells from the experiment described in B were fixed at the indicated time points and chromatin-immunoprecipitation with Whi5-V5 as bait was performed. The graph reports the total DNA yield (normalized to t1) for four independent experiments. Black bars indicate the mean of the four replicates.

Since phosphorylation of the known non-CDK sites on Whi5 did not seem to be responsible for Whi5 re-entry, we decided to further investigate the phosphorylation of CDK sites on Whi5 in response to starvation. We thus turned to a bulk population-based approach to analyze V5-tagged Whi5 by Phostag-SDS PAGE and Western blots. We synchronized cells in G1 with our previously established system [33], which is the same strain as used for constitutive Cln1 expression in Figure 6 (see also Supplementary Figure 3). As we reported previously, Whi5 leaves the nucleus approximately 35 minutes after inducing Cln1 expression by adding estradiol to the growth medium (Supplementary in [33]). We released G1-arrested cells growing on glucose minimal medium and after 45 minutes switched the cells to starvation medium. We harvested cells from the G1 arrest, 45 minutes after release immediately before starvation, and 10 and 30 minutes after the switch to starvation. These samples were analyzed by Phos-tag [40] Western blot. As reported previously [12], Whi5 becomes hyperphosphorylated as CDK activity rises (Figure 7C, lane 2). 10 minutes after starvation however, these cells begin to lose their hyperphosphorylated form of Whi5 (Figure 7C, lane 3, Supplementary Figure 5). This result suggests that Whi5 translocation is caused by *de*phosphorylation, most likely of CDK sites. Notably, we added the inducer for Cln1 expression in our starvation media, which confirms that Whi5 dephosphorylation is not dependent on cyclin repression. While our results do not exclude the possibility of additional phosphorylations on a previously uncharacterized site contributing to Whi5 re-entry, they do strongly suggest that dephosphorylation of CDK sites on Whi5 is what drives Whi5 nuclear re-entries.

We next tested whether the dephosphorylated, re-imported Whi5 also rebinds G1/S promoters. We used the same strain and set-up as described for the phosphorylation assay: Cells were arrested in G1, released by inducing Cln1 expression, and starved 45 minutes after induction. Chromatin-immunoprecipitation was then performed from these samples with the tagged Whi5. Unfortunately, we were technically not able to reliably quantify re-binding to individual promoters. However, we could reproducibly show that Whi5 pulls down less DNA after the release than in the G1 arrest. Once the released cells are starved, the amount of DNA bound to Whi5 increases again. We conclude that after starvation, Whi5 is dephosphorylated, re-enters the nucleus and rebinds promoters to re-set the Whi5-SBF-CDK feedback-loop, thus preventing the G1/S transition and re-establishing a pre-Start state.

However, these results did not tell us what the initial target of nutrient signaling is. It is possible that the specific activation of a phosphatase causes Whi5-dephosphorylation and interruption of the feedback loop. However, we are not aware of a phosphatase that is strongly upregulated upon starvation and could target Whi5. The only previously reported phosphatase targeting Whi5 in G1 is PP2A-Cdc55 [19]. The deletion mutant of Cdc55 is very sick with irregular cell cycle progression, and therefore Whi5 behavior could not be quantified reliably. However, those cells that did manage to pass Start, also showed Whi5 re-entries, indicating that Cdc55 is not essential for Whi5 dephosphorylation under starvation. We therefore consider it more likely that dephosphorylation of Whi5 is driven by an inhibition of Cdk1 kinase activity through a yet unknown mechanism, rather than by specific activity of a phosphatase.

## Discussion

In this work, we analyzed the starvation response of thousands of post-Start cells. In agreement with recent studies, we show that cells can delay or arrest their cell cycle in any cell cycle stage [21, 23]. Stable arrests are achieved mainly in G1 and G2 (as judged from budding state and histone-TFP intensity). We set out to understand how these stable arrests are achieved and found that the cell cycle inhibitor Whi5 can re-enter the nucleus in post-Start cells. We showed that Whi5 re-entries can be classified as fast and slow re-entries, which likely corresponds to cells before and after entering S-phase. Since it is unclear if Whi5 has any function in the later cell cycle, we focused on those cells re-importing Whi5 with a steep slope, which are likely to be cells before replication. In the time window of 20 minutes (which is approximately 20% of the total time between Start and cytokinesis) over 70 % of all cells respond to carbon starvation by re-importing Whi5.

We provide at least three lines of evidence that these cells are indeed reversing CDK activation and are “taking back” their commitment decision: Firstly, most cells that re-import Whi5 upon starvation become sensitive to mating pheromone again (Figure 5), which you only expect from pre-Start cells. Secondly, upon glucose replenishment these cells produce another peak of Cln2 expression, just like cells that have no history of passing Start (Figure 5). Thirdly, Whi5 re-associates with DNA (Figure 7), thus likely inhibiting expression of G1/S genes again. Therefore, it seems highly likely that cells are indeed reversing Start (as defined in the textbooks). We note that Whi5 re-entries have been previously mentioned in various contexts [21, 25, 27, 28], although nobody had mechanistically studied their cause or consequence. This shows that this is not an observation limited to our laboratory or to the strain investigated.

Are all cells that passed and reversed Start really identical to “normal” pre-Start G1 cells after starvation? Several observations suggest that while cells reverse Cdk1 activation, they may retain some “memory” of passing Start. While many of these cells start shmooing in response to mating pheromone, the fraction of cells deciding to do so is 30% lower than the fraction in “normal” G1 cells at the tested concentration. Also, cells with tiny buds that have not yet started replicating, can reverse (and later re-activate) Cln1/2-CDK activity, but they do not shmoo, and continue budding at the same site when re-entering the cell cycle after glucose replenishment. We also note that cells with Whi5 re-entries arrested with very different Sic1 concentrations. Those cells that had not started degrading Sic1, stabilized the protein and arrested with high Sic1 levels. Other cells that were starved after they had degraded Sic1 arrested in a low Sic1 state and did not express any more Sic1 while in the G1 arrest. This may be relevant for the next cell cycle following nutrient upshift. Further molecular and biochemical studies are needed to fully understand the consequences of Whi5 re-entry and Start reversal after starvation (and possible other perturbations).

While we were not able to fully unveil the mechanism that translates nutrient sensing into interruption of the positive feedback loop and Start reversal, we can make several relevant conclusions: We first analyzed a series of transcriptional regulators that are known to be important for stationary phase and quiescence. However, none of these were essential to the acute starvation response of Whi5. In fact, a transcriptional mechanism seems unlikely, because overexpression of Cln1 did not prevent or delay re-entries. We therefore turned away from transcription factors and searched for mechanisms directly acting on Whi5 and Cdk1. We showed that the mechanism of Whi5 re-import is likely the dephosphorylation of the CDK1 sites, and not due to a regulatory phosphorylation on one of the previously reported [12] non-CDK sites. However, it is still possible that one of the many uncharacterized sites that have been picked up in mass spectrometry experiments (see e. g. biogrid.org [41] entries) could contribute to Whi5’s response to starvation.

The observed dephosphorylation of Whi5 CDK sites could be caused by the upregulation of a starvation induced phosphatase. However, we did not find any evidence pointing in that direction. We therefore suggest that the cyclin-CDK complex itself is the target of starvation signaling. However, neither Sic1 nor Cip1, the two inhibitors of CDK at the G1/S transition, appear to be essential for Start reversal. Other plausible mechanisms are, for example, an inhibitory post-translational modification of CDK or the cyclins, or the selective degradation of Start components [42], which we have not been able to systematically investigate within the scope of this study.

If Whi5 export and CDK activation do not define the final commitment point, then what is the point-of-no-return? Common sense dictates that cells should irreversibly commit to the cell cycle before attempting to replicate their DNA. Although we lack a good read-out for replication initiation in budding yeast, our data on histone fluorescence intensity indeed suggests that all cells with fast Whi5 re-entries are pre-replication. In mammalian cells, APC inactivation has been suggested as the point-of-no-return [43]. In yeast cells, APC-Cdh1 inactivation was recently placed 12+/- 3 minutes after Whi5 exit using an APC substrate fragment as a sensor [44]. This time window is shorter than the ~20 minutes observed for Whi5 re-entries in our experiments, but this difference could also be due to differences in strains and nutrient conditions. So, the role of the APC in commitment to S-phase in yeast warrants further exploration.

The next step will be to find the missing pieces leading from Whi5 exit to irreversible commitment. As introduced above, the widely accepted textbook model states that the point-of-no-return in the cell cycle is “Start” in yeast and the “Restriction Point” in mammalian cells [6–8]. However, in mammalian cells, the notion that the restriction point is the universal cell cycle commitment point has been strongly challenged. Several studies suggested that there are at least two different commitment points: The restriction point for hormone and growth factor signaling, and a later point vis-à-vis nutrient and stress signals [8, 43, 45–47]. However, a complete and coherent model of all steps leading to cell cycle commitment in mammalian cells is still lacking. We now show that also in yeast commitment is a multi-step process. Thus, our yeast experiments provide an excellent foundation to quantitatively and mechanistically study eukaryotic cell cycle commitment in a simple, tractable model.

## Methods

### Strain Construction and Cultivation

All strains were haploid W303 derivatives and were prototrophic except for several deletion mutants, which were partially auxotroph as indicated in Suppl. Table 1. Strains were constructed using standard PCR-based homologous recombination. Whi5 phosphorylation mutants were generated by site directed mutagenesis on a plasmid, which was introduced into a Whi5 deletion mutant. See Suppl. Table 1 for a detailed strain list with all genotypes.

For all reported experiments, yeast cells were grown in minimal medium without amino acids (1.7 g/L yeast nitrogen base without amino acids (US Biological), 5 g/l ammonium sulphate, 50 mM potassium phthalate, pH adjusted to 5 with KOH). Single amino acids were supplemented where necessary for strains and their corresponding controls (final concentrations histidine 5mg/L, leucine 120 mg/L, uracil 20 mg/L). As a carbon source, 10 g/l glucose were added, which were then replaced by 10 g/l sorbitol in the starvation medium. Cells were incubated at 30°C on an orbital shaker at 200 rpm.

Cell cycle release and starvation experiments for Figure 7 were performed as follows: Strains derived from JE611c [33, 34] were grown in a 15 ml pre-culture on glucose minimal medium containing 40 nmol/l ß-estradiol. A main culture was inoculated with OD 0.05 and grown overnight. Cells were arrested in G1 by filtering the culture and resuspending cells in estradiol-free medium. After five hours, arrest was verified by absence of budding, as observed under a light microscope. G1 cells were released into the cell cycle by addition of 200 nM ß-estradiol. For starvation experiments, released cells were filtered 45 minutes after release, washed and resuspended in sorbitol minimal medium (also supplemented with 200 nM estradiol).

### Microfluidic cultivation

In preparation for live cell imaging experiments, cells were grown in 15 ml 1 % glucose minimum medium overnight. The next morning, the cells were transferred to a fresh culture by applying 1:15 dilution and grown for at least another 5 hours until reaching log phase (OD600 between 0.4 and 0.6).

Cells were sonicated at low power for 3 s and loaded onto a commercial microfluidics system (Y04C-02 plates, CellASIC ONIX2 system, Merck). While loading the cells, the well which contained the starvation media were kept pressurized at 1.5 psi to avoid any back flow from the glucose containing wells into the glucose-free media. During growth, the glucose media was supplied with a pressure of 3 psi. The well containing the starvation media was also pressurized with 0.5 psi to avoid backflow. Cells were grown inside the microfluidic chamber for the duration of at least 2 cell cycles before starting imaging. The temperature was kept constant at 30 °C using an incubator chamber surrounding the imaging system (Okolab Cage Incubator, Okolab USA INC, San Bruno, CA).

The mating pheromone experiments were performed using the same setup, except that 500 nM α-factor were added to the glucose media supplied after the starvation period. To prevent α-factor from adhering to the cell wall, 20 μg/ml casein was added. For the growth of JE616, a final concentration of 1 μM beta-estradiol was added to both glucose and sorbitol minimal media.

### Microscopy

All live-cell imaging experiments were performed on a Nikon Ti2 inverted epifluorescence microscope (Nikon Instruments, Japan) with a Lumencor SPECTRA X light engine (Lumencor, Beaverton, USA), a Photometrics Prime 95 (Teledyne Photometrics, USA) back-illuminated sCMOS camera. The system was programmed and controlled by the Nikon software NIS Elements. Focus was maintained using the Nikon “Perfect Focus System”. The time-lapse images were taken using a Nikon PlanApo oil-immersion 60X objective (NA=1.4) with a frequency of 3 minutes (2 minutes for experiments using the Cln2-dPSTR construct). See Suppl. Table 2 for optical filters and Suppl. Table 3 for exposure settings. For all fluorophores and tagged proteins, we checked for absence of phototoxicity by comparing growth rates at varying exposures, frame rates and in non-exposed cells. Empty strains and single-fluorophore expressing cells were used to control for autofluorescence and bleed-through, respectively.

### Image Analysis and Data Processing

Cell segmentation, tracking, fluorescence quantification and nuclear detection were performed using a custom-built Matlab script adapted from [48, 49]. All code used in this manuscript is available from the authors at request. The average autofluorescence per area was determined using a strain without fluorophores and subtracted from the recorded fluorescence intensity. For protein concentrations during cell cycle progression, total (nuclear+cytoplasmic) fluorescence intensity divided by total area was used. The segmentation of the nucleus was achieved by applying a two-dimensional Gaussian fit to the brightest pixels as described in [17]. The accuracy of this fitting method for Whi5 based nuclear detection was verified using a fluorophore-labelled histone as a nuclear marker. Start was defined as the point where 50% of the Whi5 had exited the nucleus [17]. Whi5 re-entries during starvation were defined as re-importing Whi5 back into the nucleus after exporting at least 50 % of nuclear Whi5. We note that none of our results qualitatively depend on the precise definition of Start. We verified for the wildtype that results were nearly identical when Start was defined as complete Whi5 export. When calculating Whi5 re-entry slopes, nuclear Whi5 intensity per area was scaled between the minimal and maximal intensity values of each cell. A linear regression was then fit to first 30 minutes of the increase.

For visualization purposes only, fluorescent images shown in figures were denoised by setting the lowest 5 % intensities of pixels to zero, and then rescaling the resulting images to the full range of the grey-scale. All data analysis was performed on raw 8-bit tiff images, exported from NIS-Elements software.

### Phostag SDS PAGE Western Blot

10 ml of cell culture were harvested by centrifugation, frozen in liquid nitrogen and stored at −80 °C. Cells were lysed by bead beating in urea buffer (20 mM Tris-HCl pH 7.5, 2 M Thiourea, 7 M Urea, 65 mM CHAPS, 65 mM DTT) supplemented with 1x EDTA-free protease and phosphatase inhibitor cocktails (GoldBio). 2-3 μg of total protein were loaded onto 10% SDS-polyacrylamide (29:1 Bio-Rad) gels with 10 μM phostag reagent (FUJIFILM Wako Chemicals) and 20 μM Mn2Cl. Gels were run in SDS buffer for approximately two hours at 15 mA on ice. Gels were washed with 10 mM EDTA and then blotted on a dry blotting transfer system (iBlot2, Invitrogen). Whi5-V5 was detected with a commercial Anti-V5 mouse antibody (Bio-Rad, MCA1360) and an Anti-Mouse HRP Conjugate (Promega, W4021). Luminescence was imaged on a Licor Odyssey FC.

### ChIP

The ChIP protocol was adapted from [50]. Cell cycle release and starvation experiments were performed as described above. For starvation, released cells were filtered after 45 minutes, washed and resuspended in a sorbitol minimal medium, supplemented with 200 nM estradiol. Cross-links between DNA-protein were introduced by incubating cells with 1% formaldehyde for 20 minutes at room temperature. 125 mM of glycine was added and incubated for 5 minutes to stop crosslinking. Cells were washed three times with ice-cold TBS and cell pellets were frozen. Frozen cell pellet was resuspended in lysis buffer (50mM HEPES pH 7.5, 140mM NaCl, 1% Triton, 0.1% Na Deoxycholate, 1mM EDTA and Protease Inhibitors). Cells were broken with zirconia beads 4 × 60 seconds at maximum speed in BeadBug homogenizer (Benchmark Scientific). After 15 minutes of centrifugation at maximum speed, the supernatant was discarded and the pellet (chromatin fraction) was resuspended in 400 μL of lysis buffer. The DNA was fragmented between 500 to 1000 bp by sonication with Branson Sonifier 250. After clarification, the sonicated chromatin fraction was subjected to overnight immunoprecipitation with 80 μL anti-FLAG beads (Sigma, A2220). The beads were then washed 1X with lysis buffer and 5X with wash buffer (10mM Tris-HCl pH8.0, 250mM LiCl, 0.75% NP-40, 0.75% Na Deoxycholate, 1mM EDTA). Protein-DNA complex was eluted with 50 μL ChIP elution buffer (50mM Tris-HCl pH8.0, 10mM EDTA, 1% SDS) and crosslinking was reversed by incubating the eluate overnight at 65°C. DNA was purified using QIAGEN PCR purification kit (Cat. No. 28104). 1 μl of the sample was loaded on an NP80 Spectrophotometer (Implen GmbH, Germany) and the concentration determined by absorbance at 260 nm. To compare the ChIP-DNA yield between replicates, the measured DNA concentrations were normalized to the concentrations from the arrested cells of the same experiment.

## Acknowledgements

We thank the Linda Breeden, Kurt Schmoller, Jan Skotheim, Serge Pelet, and Francesc Posas labs for strains and plasmids. We gratefully acknowledge Katja Kleemann and Stefanie Dommel for technical support. JCE acknowledges funding by the Institutional Strategy of the University of Tübingen (Deutsche Forschungsgemeinschaft ZUK 63) and from the Deutsche Forschungsgemeinschaft (Project 391105827).

## Author contributions

JCE conceived and supervised the study; DI, FPS, and PM performed experiments; DI, JCE, and IN analyzed data; AD and NEW provided code and supported data analysis; JCE and DI wrote the manuscript.

## Movies

Movie 1: **Example cell showing a Whi5 re-entry**. Cells expressing Whi5-mCherry and a Cln2-promoter-dPSTR construct (see also Supplementary Figure 1) were grown on glucose minimal medium and then switched to starvation medium as indicated by the blue background.

Movie 2: **Example cell with a Whi5 re-entry and shmoo**. Cells were starved (as indicated by blue background) and then exposed to alpha-factor when glucose was replenished.

## References

1. Smets, B., et al., Life in the midst of scarcity: adaptations to nutrient availability in Saccharomyces cerevisiae. Curr Genet, 2010. 56(1): p. 1–32.

2. Qu, Y., et al., Cell Cycle Inhibitor Whi5 Records Environmental Information to Coordinate Growth and Division in Yeast. Cell Rep, 2019. 29(4): p. 987–994 e5.

3. Broach, J.R., Nutritional control of growth and development in yeast. Genetics, 2012. 192(1): p. 73–105.

4. Ewald, J.C., How yeast coordinates metabolism, growth and division. Current Opinion in Microbiology, 2018. 45: p. 1–7.

5. Turner, J.J., J.C. Ewald, and J.M. Skotheim, Cell size control in yeast. Curr Biol, 2012. 22(9): p. R350–9.

6. Johnson, A. and J.M. Skotheim, Start and the restriction point. Curr Opin Cell Biol, 2013. 25(6): p. 717–23.

7. Morgan, D.O., ed. The Cell Cycle: Principles of Control. Primers in Biology 2007, New Science Press: London.

8. Pennycook, B.R. and A.R. Barr, Restriction point regulation at the crossroads between quiescence and cell proliferation. FEBS Lett, 2020.

9. Cross, F.R., N.E. Buchler, and J.M. Skotheim, Evolution of networks and sequences in eukaryotic cell cycle control. Philos Trans R Soc Lond B Biol Sci, 2011. 366(1584): p. 3532–44.

10. de Bruin, R.A., et al., Cln3 activates G1-specific transcription via phosphorylation of the SBF bound repressor Whi5. Cell, 2004. 117(7): p. 887–98.

11. Schmoller, K.M., et al., Dilution of the cell cycle inhibitor Whi5 controls budding-yeast cell size. Nature, 2015. 526(7572): p. 268–72.

12. Wagner, M.V., et al., Whi5 regulation by site specific CDK-phosphorylation in Saccharomyces cerevisiae. PLoS One, 2009. 4(1): p. e4300.

13. Kõivomägi, M., et al., Localized phosphorylation of RNA Polymerase II by G1 cyclin-Cdk promotes cell cycle entry. bioRxiv, 2021: p. 2021.03.25.436872.

14. Costanzo, M., et al., CDK activity antagonizes Whi5, an inhibitor of G1/S transcription in yeast. Cell, 2004. 117(7): p. 899–913.

15. Skotheim, J.M., et al., Positive feedback of G1 cyclins ensures coherent cell cycle entry. Nature, 2008. 454(7202): p. 291–6.

16. Charvin, G., et al., Origin of irreversibility of cell cycle start in budding yeast. PLoS Biol, 2010. 8(1): p. e1000284.

17. Doncic, A., M. Falleur-Fettig, and J.M. Skotheim, Distinct interactions select and maintain a specific cell fate. Mol Cell, 2011. 43(4): p. 528–39.

18. Qu, Y.M., et al., Cell Cycle Inhibitor Whi5 Records Environmental Information to Coordinate Growth and Division in Yeast. Cell Reports, 2019. 29(4): p. 987–+.

19. Talarek, N., E. Gueydon, and E. Schwob, Homeostatic control of START through negative feedback between Cln3-Cdk1 and Rim15/Greatwall kinase in budding yeast. Elife, 2017. 6.

20. Duch, A., et al., Multiple signaling kinases target Mrc1 to prevent genomic instability triggered by transcription-replication conflicts. Nat Commun, 2018. 9(1): p. 379.

21. Wood, N.E., et al., Nutrient Signaling, Stress Response, and Inter-organelle Communication Are Non-canonical Determinants of Cell Fate. Cell Rep, 2020. 33(9): p. 108446.

22. Laporte, D., et al., Metabolic status rather than cell cycle signals control quiescence entry and exit. J Cell Biol, 2011. 192(6): p. 949–57.

23. Argüello-Miranda, O., et al., Cell cycle-independent integration of stress signals promotes Non-G1/G0 quiescence entry. bioRxiv, 2021: p. 2021.03.13.434817.

24. Bertoli, C., J.M. Skotheim, and R.A. de Bruin, Control of cell cycle transcription during G1 and S phases. Nat Rev Mol Cell Biol, 2013. 14(8): p. 518–28.

25. Liu, X., et al., Reliable cell cycle commitment in budding yeast is ensured by signal integration. Elife, 2015. 4.

26. Aymoz, D., et al., Real-time quantification of protein expression at the single-cell level via dynamic protein synthesis translocation reporters. Nat Commun, 2016. 7: p. 11304.

27. Litsios, A., Metabolic-rate dependent cell cycle entry and progression in Saccharomyces cerevisiae. 2017, University of Groningen: Groningen. p. 148.

28. Litsios, A., Metabolic-rate dependent cell cycle entry and progression in Saccharomyces cerevisiae. 2017, University of Groningen: University of Groningen.

29. Pirincci Ercan, D., et al., Budding yeast relies on G1 cyclin specificity to couple cell cycle progression with morphogenetic development. Sci Adv, 2021. 7(23).

30. Miles, S., et al., Xbp1 directs global repression of budding yeast transcription during the transition to quiescence and is important for the longevity and reversibility of the quiescent state. PLoS Genet, 2013. 9(10): p. e1003854.

31. Monteiro, P.T., et al., YEASTRACT plus : a portal for cross-species comparative genomics of transcription regulation in yeasts. Nucleic Acids Research, 2020. 48(D1): p. D642–D649.

32. Miles, S., et al., Msa1 and Msa2 Modulate G1-Specific Transcription to Promote G1 Arrest and the Transition to Quiescence in Budding Yeast. PLoS Genet, 2016. 12(6): p. e1006088.

33. Ewald, J.C., et al., The yeast cyclin-dependent kinase routes carbon fluxes to fuel cell cycle progression. Molecular Cell, 2016.

34. Zhang, L., et al., Multiple Layers of Phospho-Regulation Coordinate Metabolism and the Cell Cycle in Budding Yeast. Front Cell Dev Biol, 2019. 7: p. 338.

35. Ottoz, D.S., F. Rudolf, and J. Stelling, Inducible, tightly regulated and growth condition-independent transcription factor in Saccharomyces cerevisiae. Nucleic Acids Res, 2014.

36. Escote, X., et al., Hog1 mediates cell-cycle arrest in G1 phase by the dual targeting of Sic1. Nature Cell Biology, 2004. 6(10): p. 997–+.

37. Adrover, M.A., et al., Time-Dependent Quantitative Multicomponent Control of the G(1)-S Network by the Stress-Activated Protein Kinase Hog1 upon Osmostress. Science Signaling, 2011. 4(192).

38. Chang, Y.L., et al., Yeast Cip1 is activated by environmental stress to inhibit Cdk1-G1 cyclins via Mcm1 and Msn2/4. Nature Communications, 2017. 8.

39. Gonzalez-Novo, A., et al., Hog1 Targets Whi5 and Msa1 Transcription Factors To Downregulate Cyclin Expression upon Stress. Molecular and Cellular Biology, 2015. 35(9): p. 1606–1618.

40. Kinoshita, E., et al., Phosphate-binding tag, a new tool to visualize phosphorylated proteins. Mol Cell Proteomics, 2006. 5(4): p. 749–57.

41. Oughtred, R., et al., The BioGRID database: A comprehensive biomedical resource of curated protein, genetic, and chemical interactions. Protein Science, 2021. 30(1): p. 187–200.

42. Morshed, S., et al., TORC1 regulates G1/S transition and cell proliferation via the E2F homologs MBF and SBF in yeast. Biochem Biophys Res Commun, 2020. 529(3): p. 846–853.

43. Cappell, S.D., et al., Irreversible APC(Cdh1) Inactivation Underlies the Point of No Return for Cell-Cycle Entry. Cell, 2016. 166(1): p. 167–80.

44. Ondracka, A., J.A. Robbins, and F.R. Cross, An APC/C-Cdh1 Biosensor Reveals the Dynamics of Cdh1 Inactivation at the G1/S Transition. PLoS One, 2016. 11(7): p. e0159166.

45. Yang, H.W., et al., Stress-mediated exit to quiescence restricted by increasing persistence in CDK4/6 activation. Elife, 2020. 9.

46. Patel, D., et al., A Late G1 Lipid Checkpoint That Is Dysregulated in Clear Cell Renal Carcinoma Cells. J Biol Chem, 2017. 292(3): p. 936–944.

47. Foster, D.A., et al., Regulation of G1 Cell Cycle Progression: Distinguishing the Restriction Point from a Nutrient-Sensing Cell Growth Checkpoint(s). Genes Cancer, 2010. 1(11): p. 1124–31.

48. Wood, N.E. and A. Doncic, A fully-automated, robust, and versatile algorithm for long-term budding yeast segmentation and tracking. PLoS One, 2019. 14(3): p. e0206395.

49. Doncic, A., et al., An algorithm to automate yeast segmentation and tracking. PLoS One, 2013. 8(3): p. e57970.

50. Flick, K.M., et al., Grr1-dependent inactivation of Mth1 mediates glucose-induced dissociation of Rgt1 from HXT gene promoters. Molecular Biology of the Cell, 2003. 14(8): p. 3230–3241.

