## Supplementary Material for "When Yeast Cells Change their Mind: Cell Cycle “Start” is Reversible under Starvation"

**Supplementary Tables:**

Table 1. Yeast strains used in this study

Table 2. Optical filters for microscopy

Table 3. Exposure times and intensities

**Supplementary Figures:**

Figure 1. Constructs to analyze Cln2 promoter activity

Figure 2. Msn2 localization upon starvation

Figure 3. The estradiol promoter is active under starvation

Figure 4. Sic1 concentrations at Whi5 re-entry

Figure 5. Phostag Western Blot (replicate of Figure 7C)

**Supplementary Table 1**: Yeast strains used in this study. All strains are W303 derivatives.

| Name | Figures | Genotype | Source |
|---|---|---|---|
| FS042 | Figure 5<br>Figure 6A | MAT a, ADE2, TRP1, LEU2, URA3, HIS | This study |
| DI002 | Figure 1B,C<br>Figure 2<br>Figure 3A,B,C<br>Figure 5 D,E,F<br>Figure 6A | MAT a, Htb2-mTFP-Nat, Whi5-mCherry-KanMX,<br>Sic1-mNeongreen-hph, ADE2, LEU2, HIS, TRP1, URA3 | This Study |
| DI028 | Figure 1C<br>Figure 2<br>Figure 3A,B,C<br>Figure 4<br>Figure 5A,B,C<br>Figure 6A | MAT a, Whi5-mCherry-KanMX, ura3::Cln2-promoter-mNeongreen-URA3, ADE2, TRP1, LEU2, | This Study |
| DI011 | Figure1C Figure 2<br>Figure 3A,B,C<br>Figure 4<br>Figure 6A | MAT α, ADE22, Whi5-mNeongreen-Ura3 | This Study |
| FS034 | Figure 2 | MAT a, Whi5-mCherry-KanMX, leu2::cln2-promoter-dPSTR-mCitrine-LEU2, ADE2, TRP1, URA | Ewald Lab [1] |
| FS002 | Figure 1C<br>Figure 2<br>Figure 3<br>Figure 4<br>Figure 5 D,E,F<br>Figure 6B | MAT a, Whi5-mCherry-KanMX, Sic1-mNeongreen-hph<br>ADE2, TRP1, LEU2, URA3, HIS3 | This study |
| KCY003 | | mat α, whi5Δ::CglaTRP11, ADE2 | Kind gift of Kora-Lee Claude and Kurt Schmoller |
| DI011 | Figure 7A | MAT α, whi5::CglaTRP11, ura3::Whi5-wildtype-mNeongreen-URA, ADE2<br>LEU2 | This Study<br>(from KCY003) |
| DI041 | Figure 7A | MAT α, whi5::CglaTRP11, ura3::Whi5-7A*-mNeongreen-URA, ADE2<br>LEU2 | This Study<br>(from KCY003) |

| | | | |
|---|---|---|---|
| | | *S78, S113, S114, S149, S76, T281, S288 mutated to alanine | |
| DI031 | Figure 6A | MAT a, Whi5-mCherry-KanMX, Cip1::HphMX ADE2, TRP1, LEU2, URA3 | This Study |
| DI045 | Figure 6A | MAT α, Whi5-mCherry-KanMX, Msn2::HphMX Msn4::His, TRP1, LEU2, URA3 | This Study |
| BY6883 | Figure 6A | MAT a, Msa1::HIS3, Msa2::KanMX ADE2, TRP1, LEU2, URA3 | Breeden Lab [2] |
| DI047 | Figure 6A | MAT a, Msa1::HIS3, Msa2::KanMX Whi5-mNeongreen-Hph, TRP1, LEU2, URA3 | This Study |
| BY6602 | | MAT a, Xbp1::KanMX, ADE2, TRP1, LEU2, URA3 | Breeden Lab [2] |
| DI043 | Figure 6A | MAT a, Whi5-mNeongreen-HphMX, Xbp1::KanMX, TRP1, LEU2, URA3 | This Study (from BY6602) |
| BY6828 | | MAT a, Sic1::KanMX, ADE2, LEU2, TRP1, HIS, URA3 | Breeden Lab [2] |
| DI038 | Figure 6A | MAT a, Whi5-mNeongreen-HphMX, Sic1::KanMX ADE2, LEU2, TRP1, HIS3, URA3 | This Study (from BY6828) |
| JE616 | Figure 6A | MAT α, cln1Δ, cln2Δ, cln3::LEU2, lexOPr-Cln1-Leu2, ADE2, his3::cyc1-Pr-lexO TF-HIS3, TRP1, URA3, Whi5-GFP-NatMX | Ewald Lab [3] |
| KK079 | Figure 7 | mat α, cln1Δ, cln2Δ, cln3::LEU2, lexOPr-Cln1-Leu2, ADE2, his3::cyc1-Pr-lexO TF-HIS3, TRP1, URA, Whi5::Whi5-V5-KanMX | This Study |
| DI036 | | mat a, Whi5-mCherry-KanMX, Htb2-mTFP-NatMX, cdc55::hphMX, ADE2, TRP1, LEU2, URA3 | This Study |
| KK087 | Suppl.3 Figure | mat α, ADE2, TRP1, LEU2, his3::LexO transcription factor- HIS3, ura3:: LexA promoter-mKO-URA3(estradiol inducible mKO) | This Study |

**Supplementary Table 2:** Optical filters for microscopy

Filters used for imaging on a Nikon Ti2-E epifluorescence microscope.
All described filters are manufactured by Chroma and purchased from AHF.

| Fluorophore | LED Wave-length | Filter Set | Excitation Filter | Dichroic | Emission Filter |
|---|---|---|---|---|---|
| mTFP | 475 nm | TFP ET Filter Set | ET445/30x | T470lpxr, Di 25 mm x 36 mm | ZET488/10x |
| Neongreen | 513 nm | YFP ET Filter Set | ET500/20x | T515lp, Di 25 mm x 36 mm | ET535/30m |
| mCitrine | 513 nm | YFP ET Filter Set | ET500/20x | T515lp, Di 25 mm x 36 mm | ET535/30m |
| mCherry | 575 nm | CFP/YFP/mCherry/ Cy7 QUAD LED ET | Quadband Exciter ET/422-449/496-517/566-588/709-752 | Quadband Beamsplitter (89402bs) | Quadband Emitter ET/463-484/ 529-550/ 605-678/ 773-845 |

**Supplementary Table 3:** Exposure times and intensities.

| Fluorophore | Imaged Protein | Intensity | Exposure Time |
|---|---|---|---|
| mTFP | Htb2 | 10 % | 100 ms |
| Neongreen | Whi5, Sic1, Cln2-Promoter, Msn4 | 15 % - 30 % | 200 ms - 300 ms |
| mCitrine | Cln2-promoter-dPSTR | 30 % | 300 ms |
| mCherry | Whi5 | 40 % - 60 % | 300 ms - 600 ms |

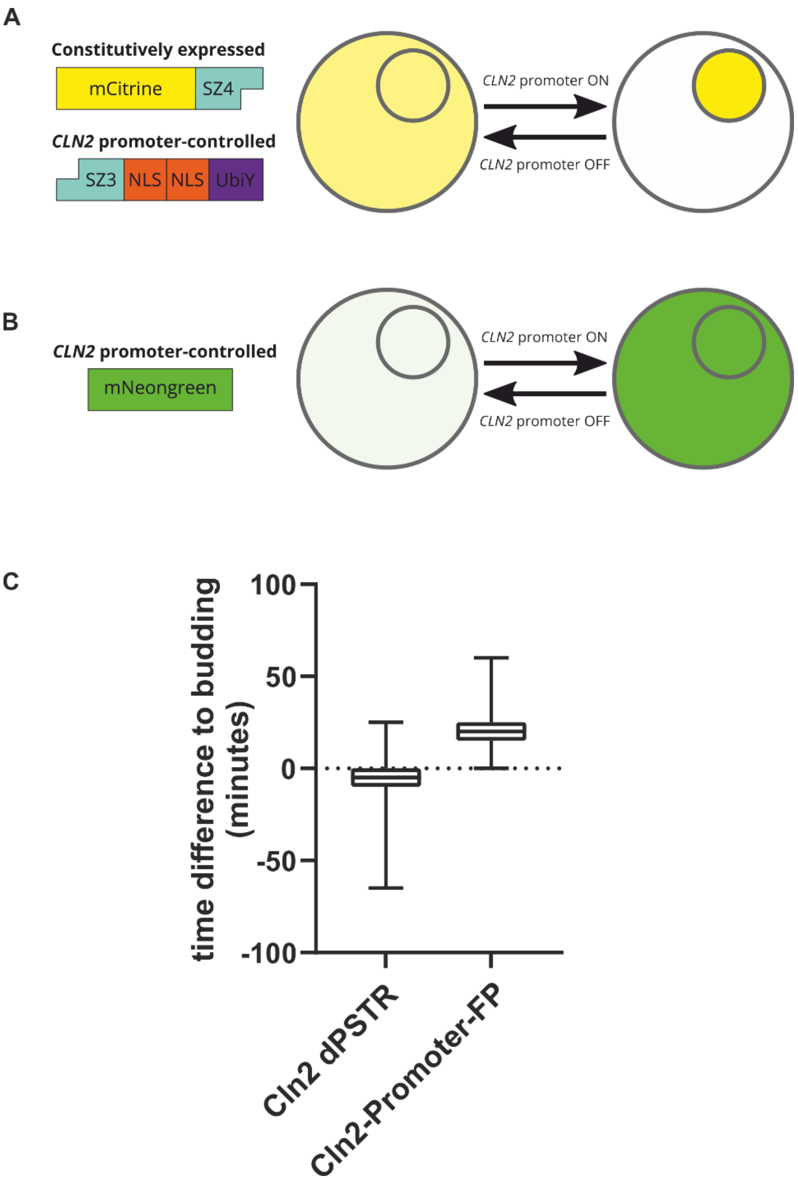


**Supplementary Figure 1**. **Cln2 promoter reporters used in this study**. **A.** The dynamic protein synthesis translocation reporters (dPSTR) developed by the Pelet lab [4] circumvent the time delay in expression analysis caused by fluorescent protein maturation. A fluorescent protein is constitutively expressed, while a small protein containing an interaction domain and a nuclear localization sequence is under control of the promoter of interest. We used this in Figure 2 to monitor the precise timing of Cln2 promoter activation in cells that passed Start before encountering starvation. Since we observed a decrease in total fluorescence during starvation from this construct, we used another construct to monitor Cln2 promoter activity after glucose re-addition. **B.** To monitor Cln2-promoter activity after starvation in Figure 5, we constructed a plasmid containing a Cln2-promoter driving Neongreen expression and integrated it at the URA3 locus. **C.** We analyzed the timing of the two reporter constructs relative to budding in unperturbed cells. The box plots indicate the time difference between budding and the peak in the fluorescent signal. Boxes indicate the median and the 25$^{th}$ and 75$^{th}$ percentile, bars indicate 5$^{th}$ and 95$^{th}$ percentile.

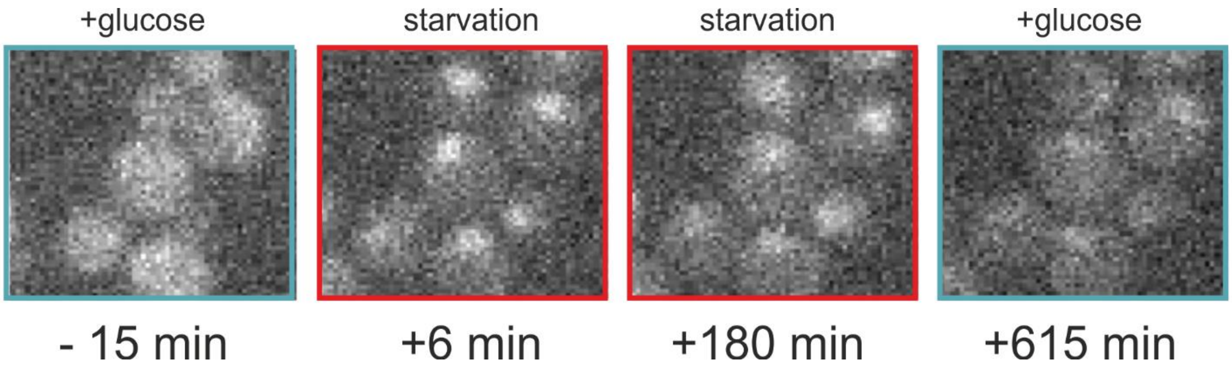


**Supplementary Figure 2: The response of Msn2 to starvation medium.** Cells expressing Msn2-Neongreen were grown for six hours on glucose minimal medium and then switched to sorbitol minimal medium at t=0 minutes. All observed cells imported Msn2 to the nucleus within 10 minutes.

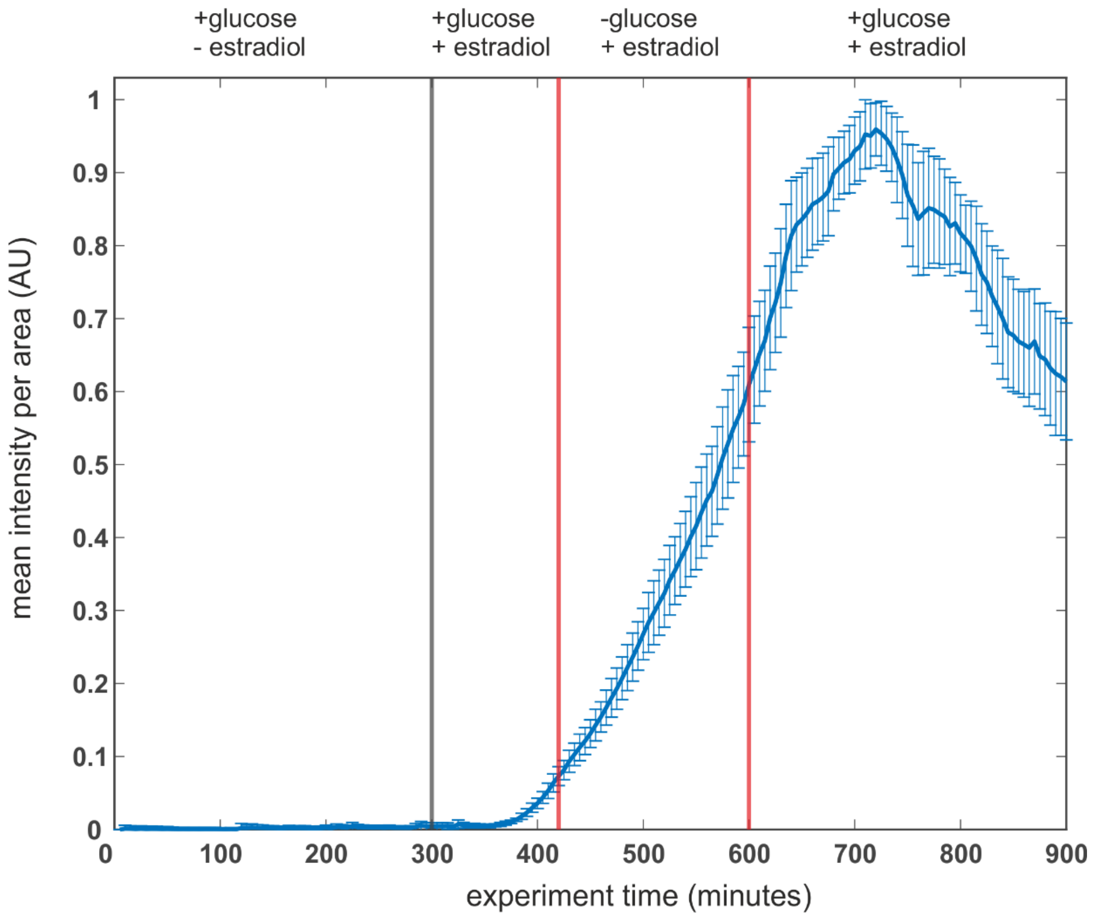


**Supplementary Figure 3: The estradiol promoter is active under starvation.** To test if the estradiol promoter [5] retains activity under starvation, we constructed a plasmid where the estradiol promoter drives expression of the fluorophore mKo. This plasmid was integrated into the ura3 locus. Cells were grown on glucose minimal medium for six hours, when mKo expression was induced by adding 1μM estradiol (the same amount as used for the Cln1 promoter). Two hours later we switched to starvation medium, but maintained the estradiol concentration. Because cells stop growing, but the estradiol promoter is active, cells continuously accumulate fluorescence signal. Shown is the mean and standard deviation of 16 cells.

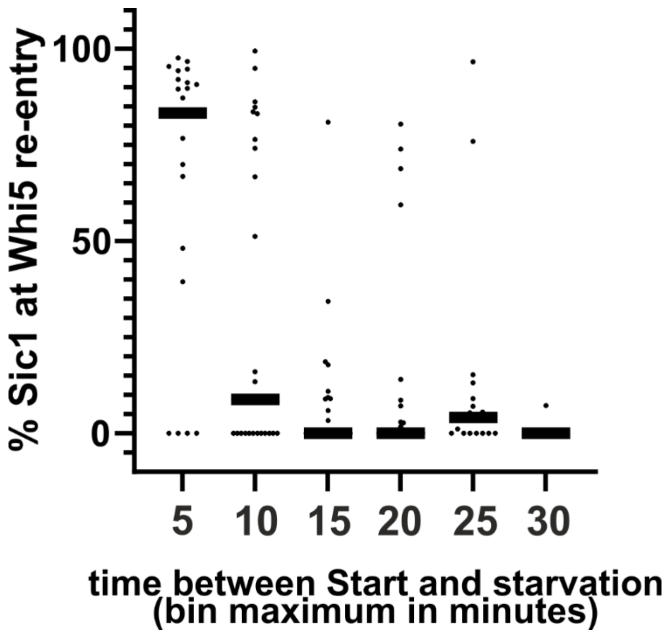


**Supplementary Figure 4: Whi5 re-entry is independent of Sic1 concentrations.** For all cells that showed fast re-entries of Whi5 (i.e. reverse Start), we determined the concentration of Sic1 (total fluorescence per area) at the beginning of Whi5 re-entry. Each dot represents an individual cell (n=149, from two biological replicates), bars represent the bin median.

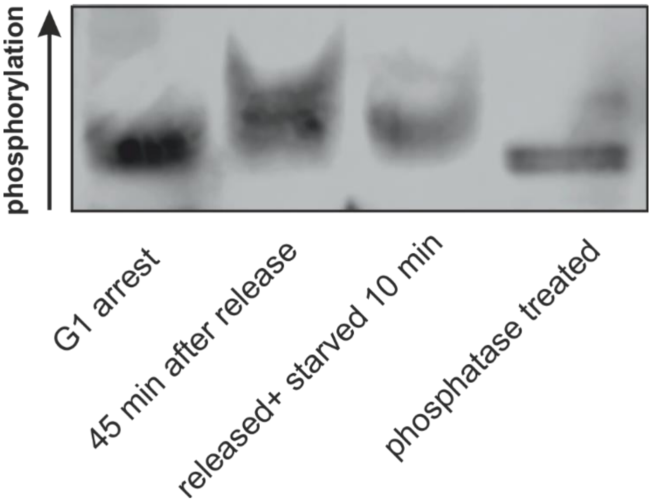


**Supplementary Figure 5. Replicate of Phostag [6] gel in Figure 7.** An independent experiment was performed as described in Figure 7B.

**Supplementary References**


1. Nadelson, I., *Sensors for monitoring cell cycle regulation during nutrient perturbations*. 2020, Eberhard Karls University of Tuebingen.
2. Miles, S., et al., *Msa1 and Msa2 Modulate G1-Specific Transcription to Promote G1 Arrest and the Transition to Quiescence in Budding Yeast.* PLoS Genet, 2016. **12**(6): p. e1006088.
3. Ewald, J.C., et al., *The yeast cyclin-dependent kinase routes carbon fluxes to fuel cell cycle progression.* Molecular Cell, 2016.
4. Aymoz, D., et al., *Real-time quantification of protein expression at the single-cell level via dynamic protein synthesis translocation reporters.* Nat Commun, 2016. **7**: p. 11304.
5. Ottoz, D.S., F. Rudolf, and J. Stelling, *Inducible, tightly regulated and growth condition-independent transcription factor in Saccharomyces cerevisiae.* Nucleic Acids Res, 2014.
6. Kinoshita, E., et al., *Phosphate-binding tag, a new tool to visualize phosphorylated proteins.* Mol Cell Proteomics, 2006. **5**(4): p. 749-57.
